# Two evolutionary distinct effectors from a nematode and virus target RanGAP1 and 2 via the WPP domain to promote disease

**DOI:** 10.1101/2021.06.24.449730

**Authors:** Octavina C. A Sukarta, Amalia Diaz-Granados, Erik J Slootweg, Hein Overmars, Casper van Schaik, Somnath Pokhare, Jan Roosien, Rikus Pomp, Abdelnaser Elashry, Geert Smant, Aska Goverse

## Abstract

The Gpa2 and Rx1 intracellular immune receptors are canonical CC-NB-LRR proteins belonging to the same *R* gene cluster in potato. Despite sharing high sequence homology, they have evolved to provide defence against unrelated pathogens. Gpa2 detects Gp-RBP-1 effectors secreted by the potato cyst nematode *Globodera pallida* whereas Rx1 recognizes the viral coat protein (CP) of Potato Virus X (PVX). How Gpa2 and Rx1 perceive their matching effectors remains unknown. Using a combination of *in planta* Co-Immunoprecipitation and cellular imaging, we show that both Gp-RBP-1 and PVX-CP physically interact with RanGAP2 and RanGAP1 in the cytoplasm of plant cells. Interestingly, this was also demonstrated for the eliciting variants of Gp-RBP-1 and PVX-CP indicating a role for RanGAP1 and RanGAP2 in pathogenicity independent from Gpa2 and Rx1 recognition. Indeed, knocking down both RanGAP homologs reduce cyst nematode and PVX infection. These findings show that RanGAP1/2 act as common host targets of evolutionary distinct effectors from two plant pathogens with different lifestyles. The involvement of RanGAP1/2 to pathogen virulence is a novel role not yet reported for these key host cell components and as such, their possible role in cyst nematode parasitism and viral pathogenicity are discussed. Moreover, from these findings a model emerges for their possible role as co-factor in pathogen recognition by the potato immune receptors Gpa2/Rx1.

## INTRODUCTION

Effective immunity hinges on the successful recognition of the invading pathogen or the damage they cause. In plants, this process is mediated by a group of cell-autonomous receptor proteins, most of which belong to the family of Nucleotide-Binding Leucine-Rich Repeats (NB-LRR) receptors (1). Plant NB-LRR immune receptors function intracellularly to detect specific pathogen-derived effector molecules. In parallel, effectors evolved to manipulate host cellular processes and/or suppress plant defence in favour of promoting virulence of the pathogen (2). Upon effector recognition, NB-LRRs trigger a suit of defence responses that can effectively suppress further infection in a process known as Effector-Triggered Immunity (ETI). This often manifests in the form of localized cell death, also known as the Hypersensitive Response (HR) (1).

Over the years, several studies have detailed how plant NB-LRRs can perceive pathogens, which have advanced our understanding of the mechanistic basis of effector recognition (reviewed in (1)). This includes a receptor-ligand model, which involves direct interaction between the NB-LRR and its matching effector. This was first documented in the rice CC-NB-LRR (CNL) Pita, for which functionality was reported to be compromised when substituting a single amino acid in its Leucine-Rich Repeat (LRR) domain. Such a substitution abolished interaction with the *Magnaporthe grisea* effector Avr-Pita (3). It appears, however, that the direct recognition model applies only to a few exceptional cases (4, 5). Instead, a majority of NB-LRRs indirectly sense pathogen-induced modifications of effector targets or their mimics (4). In this manner, it is believed that the plant can circumvent rapidly evolving pathogens by enabling a single NB-LRR to detect multiple effectors that act on a single host target. Today, a wide variety of models for pathogen detection have been described that reconcile the detection of a plethora of invading pathogens with only a limited set of NB-LRR immune receptors (1).

The CC-NB-LRRs Gpa2 and Rx1 from potato (*Solanum tuberosum* spp. *andigena*) are highly homologous immune receptors (88% identity) that mediate distinct defense responses against evolutionarily unrelated pathogens (6–8). Gpa2 confers defense against the potato cyst nematode *Globodera pallida* by inducing a hypersensitive response, disconnecting the nematode’s feeding site from the root vasculature, resulting in nematode arrest. Heterologous studies have shown, however, that Gpa2 can also trigger HR in the leaves of *Nicotiana benthamiana* upon detection of the nematode-secreted effector Gp-RBP-1 (9). In turn, Rx1 mediates immunity to Potato Virus X (PVX), a filamentous positive-sense RNA virus that infects aerial parts of *Solanaceous* plants. Upon recognition of the viral coat protein (PVX-CP), Rx1 activates a symptomless defence response referred to as extreme resistance that effectively limits infection to initially affected cells (7). Rx1 also has the capacity to induce a classical HR, when it is overexpressed or when there is an overaccumulation of PVX-CP in heterologous studies (7, 10). In contrast to the Rx1 and Gpa2 immune receptors, PVX-CP and Gp-RBP-1 effectors share no sequence nor structural similarities. It is well established that the recognition specificity of Rx1 and Gpa2 is confined to the C-terminal end of the LRR domain (11–13). Moreover, it has been demonstrated that subtle changes in amino acid residues are sufficient to evade Rx1 and Gpa2 recognition. For example, the substitution of a proline by a serine in Gp-RBP-1 prevents the activation of HR (Sacco et al 2009), whereas two amino acid substitutions in the PVX-CP compromises Rx1 recognition (7). However, the molecular mechanisms underlying effector recognition by Gpa2 and Rx1 are still unknown.

Interestingly, sequence exchange events between Rx1 and Gpa2 pinpoint to a functional bifurcation of the receptor in which recognition specificity is determined by the hypervariable LRR domain, whereas defence activation is confined to the CC-NB-ARC moiety (12). This is consistent with findings that the N-terminal moieties of plant NB-LRRs can act as scaffolds for interactions with host proteins involved in downstream signalling (14). As the CC-NB-ARC is interchangeable between Gpa2 and Rx1, this also suggests that both receptors are likely to share similar co-factors. Indeed, the conserved CC domains of both receptors form a complex with the WPP domain of the RanGTPase activating protein 2 (RanGAP2) (15, 16). RanGAP2 plays a vital role in the cell by regulating mitosis and nucleocytoplasmic transport during plant development (9, 17, 18). As such, RanGAP2 was shown to act as a co-factor by balancing the distribution of Rx1 in the nucleus and cytoplasm as well as modulating its stability (19). Although it was shown that RanGAP2 contributes to Rx1 and Gpa2-mediated defence responses, the underlying mechanisms remain unclear (15, 16).

Artificial tethering of Gp-RBP-1 to RanGAP2 in a YFP-complementation experiments showed that HR by Gpa2 in *N. benthamiana* is enhanced (9). This suggests that RanGAP2 may contribute to Gpa2-mediated immunity by facilitating Gp-RBP-1 recognition. Given that RanGAP2 is also a co-factor of Rx1, we hypothesized that RanGAP2 could also contribute to PVX-CP recognition through complex formation in plant cells. To test whether Gp-RBP-1 and PVX-CP could associate with RanGAP2 *in planta*, a combination of Co-Immunoprecipitation (Co-IP) and advanced cellular imaging studies was performed. We were able to show that RanGAP2 can indeed form protein complexes with Gp-RBP-1 and PVX-CPs *in planta* via its conserved WPP domain. We could further demonstrate that these effectors also target RanGAP1, a homolog shown previously to interact with Rx1 in a yeast-two-hybrid assay (19). Interestingly, also non-eliciting variants of PVX-CP and Gp-RBP-1 can associate with RanGAP1 and RanGAP2, suggesting a broader role for this common effector target in promoting nematode and viral pathogenicity. Indeed, knocking down either or both RanGAP homologs reduced infection by PVX in *N. benthamiana* and the cyst nematode *Heterodera schachtii* in *Arabidopsis thaliana.* These data support a model of RanGAP1/2 as a common effector target of two taxonomically distinct pathogens with different modes of action, a virus and nematode. To our knowledge, this is the first study demonstrating the effector targeting of RanGAP1/2 and their role in promoting disease in different plant parts. Possible roles of RanGAP1/2 in cyst nematode and PVX pathogenicity are discussed as well as a tripartite model for effector recognition by the immune receptors Gpa2/Rx1 that emerges from our findings.

## RESULTS

### Eliciting and non-eliciting effectors of *G. pallida* and PVX interact with full-length RanGAP1 and RanGAP2 *in planta*

To discern whether RanGAP2 and Gp-RBP-1 can form a complex *in planta*, we performed a Co-IP assay. To that end, full-length RanGAP2-GFP was co-expressed transiently in leaves of *N. benthamiana* by agroinfiltration with 8×HA-tagged versions of the Gpa2-activating or non-activating Gp-RBP-1 variants, namely D383-1 and Rook4. In plants, RanGAP2 is homologous to RanGAP1, sharing 66.2% identity at the amino acid level in *N. benthamiana*. Both protein homologs are functionally redundant where they act as activators of RanGTPase as part of the nucleocytoplasmic transport cycle (20, 21). We, therefore, sought to investigate whether RanGAP1 could also associate with Gp-RBP-1. For Co-IP, RanGAP2-GFP or RanGAP1-GFP was captured with anti-GFP (α-GFP) conjugated paramagnetic beads as bait, and the bound proteins were analysed by immunoblotting (**Fig. 1A**). Our data show that there were no changes in protein stability of RanGAP1-GFP, RanGAP2-GFP or the Gp-RBP-1 effectors when co-expressed. Both D383-1-8×HA and Rook4-8×HA effectors specifically co-immunoprecipitated with both RanGAP1-GFP and RanGAP2-GFP. Interestingly, these *G. pallida* effectors co-purified more in the eluate in combination with the RanGAP2 homolog compared to RanGAP1. Moreover, stronger band intensity for the non-eliciting Rook4 variant compared to D383-1 was consistently observed after Co-IP when RanGAP1 and RanGAP2 was used as bait. This suggests that eliciting and non-eliciting Gp-RBP-1 effectors may differ in their binding affinities for RanGAP2 and RanGAP1. Combined, our results demonstrate that both eliciting and non-eliciting Gp-RBP-1 effectors can form complexes with both RanGAP homologs *in planta*.

**Fig. 1.**
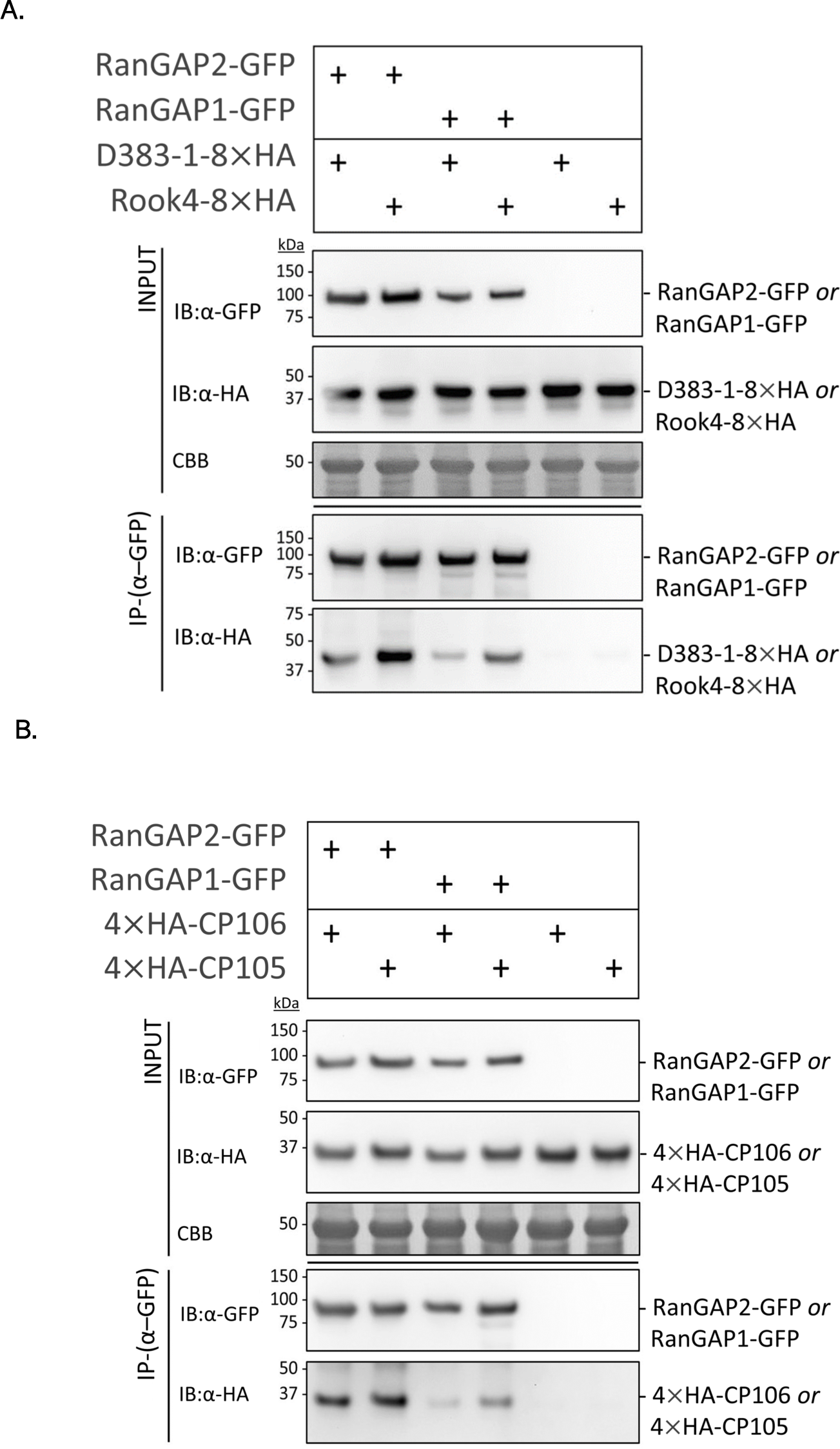
Gp-RBP-1 of *G. pallida* and CP of PVX associate with RanGAP1 and RanGAP2 *in planta*. Co-immunoprecipitation of full-length RanGAP1-GFP or RanGAP2-GFP as bait and HA-tagged Gp-RBP-1 (**A**) or PVX-CP (**B**) effector proteins. *A. tumefaciens* harbouring constructs for the pull-downs were co-expressed in *N. benthamiana* leaves and harvested at 2 dpi. “+” indicates the presence of a construct in the infiltration combination. The soluble extract was used for Co-IP studies using α-GFP conjugated beads to precipitate the bait. The immunoblots (IB) with α-GFP and α-HA antibodies of the input material are shown in the top half of the image and the results of Co-IP in the two bottom panels of the figure. Coomassie brilliant blue (CBB) stained blots serve as a loading control for the input material. Data shown are representative of three independent repeats.

As RanGAP2 was initially found to be a co-factor of Rx1, we expanded our Co-IP studies to explore whether RanGAP1 and/or RanGAP2 can also form a complex with PVX-CP (15, 16). Full-length RanGAP1-GFP and RanGAP2-GFP were used as bait to pull-down 4×HA-tagged versions of the coat proteins from the non-eliciting UK3 (CP106) and eliciting HB (CP105) PVX strains. In line with the interaction data for Gp-RBP-1, we did not observe changes in protein stabilities of RanGAP1-GFP, RanGAP2-GFP, CP105-4×HA, or CP106-4×HA upon co-expression (**Fig. 1B**). Both CP106-4×HA and CP105-4×HA variants co-immunoprecipitated with RanGAP2-GFP, whereas no detectable amounts were pulled down with the α-GFP beads alone. Our findings, therefore, reveal that also eliciting and non-eliciting variants of PVX-CP can associate with both RanGAP homologs *in planta* like Gp-RBP-1. RanGAP1 and RanGAP2 are, therefore, common interactors of structurally divergent effector types from taxonomically unrelated pathogens with distinct lifestyles. Notably, both viral coat protein variants also consistently co-purified in lower quantities when RanGAP1-GFP was used as bait compared to RanGAP2-GFP like observed for Gp-RBP-1.

### RanGAP1 and RanGAP2 is required for virulence by nematodes and viruses

Given that RanGAP1 and RanGAP2 can interact with both the eliciting and non-eliciting variants of PVX-CP and Gp-RBP-1, we hypothesized that plant RanGAPs could fulfil a broader role beyond functioning as a co-factor in pathogen recognition. For instance, the effector targeting of host proteins is known to be directly used by pathogens to promote virulence (22). Thus, we investigated whether RanGAP1 and/or RanGAP2 can contribute to plant susceptibility to nematode and/or viral infections. To investigate the function of RanGAP1 and RanGAP2 in PVX infection, we performed Tobacco Rattle Virus-based Virus-Induced Gene Silencing (TRV-VIGS) in *N. benthamiana* using constructs described in (15, 19). Twenty-one days after inoculation with TRV, leaves *N. benthamiana* plants were infiltrated with agrobacteria harbouring amplicons of PVX-106 (non-eliciting) or PVX-105 (eliciting). Viral levels were quantified in the infiltrated zones within 1-5 dpi by DAS-ELISA. Our data showed that when RanGAP2 or RanGAP1 is silenced, significantly less viral accumulation occurs compared to the TRV:GFP control irrespective of the viral strain at 3 dpi but not at 5 dpi (**Supplementals Fig. S5**) (**Fig. 2B**). Simultaneously silencing RanGAP1 and RanGAP2 by VIGS results in greater suppression of both virus. Our TRV-VIGS data, therefore, illustrate that both RanGAP2 and RanGAP1 contribute to PVX virulence in *N. benthamiana*, only during the early stages of viral infection.

**Fig. 2.**
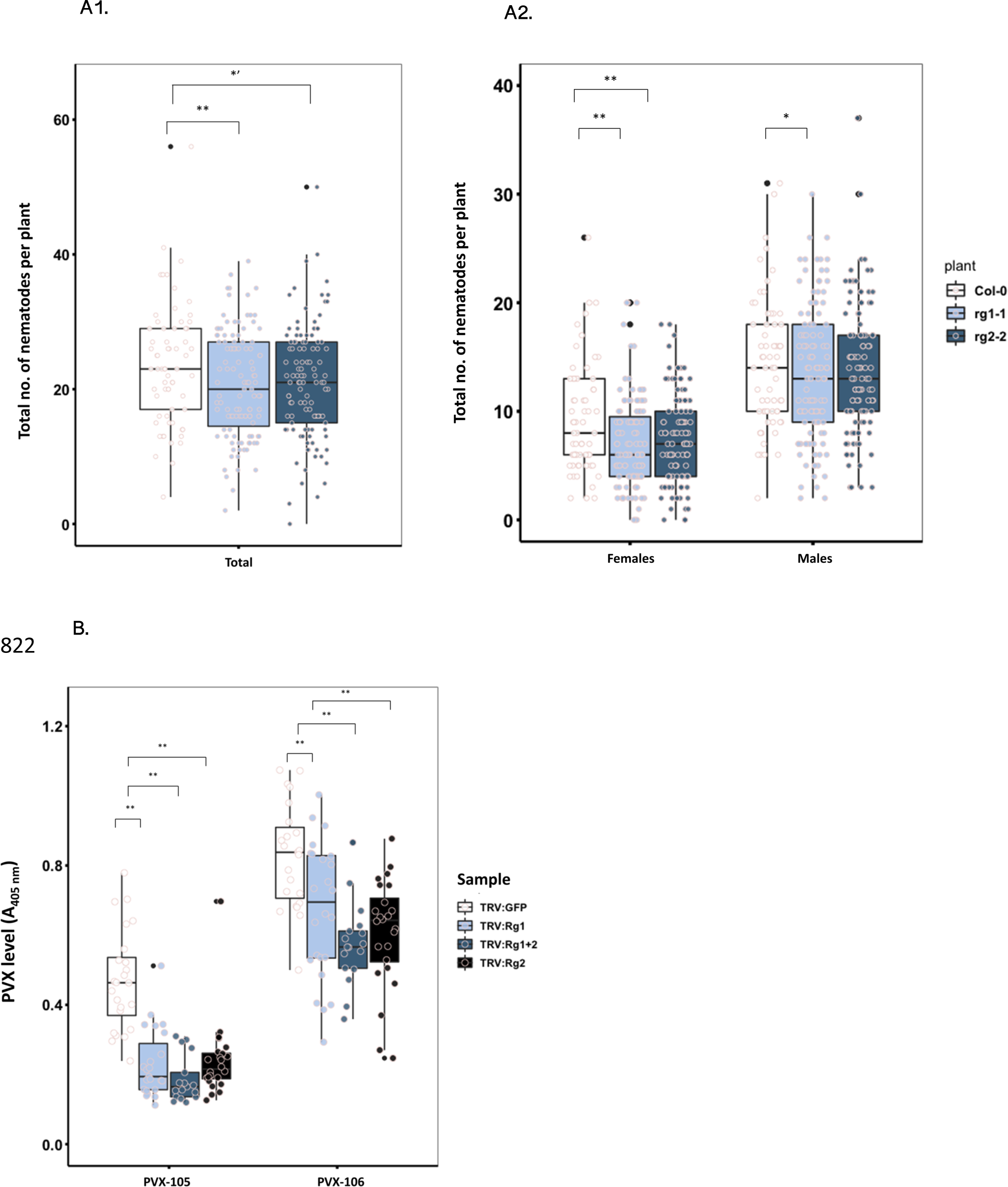
RanGAP1/2 contributes to pathogenicity of cyst nematodes in *A. thaliana* and PVX in *N. benthamiana*. **A).** The total (**A1**) and average number of female and male (**A2**) nematodes per plant in *A. thaliana* roots after 2 weeks of infection. Boxes indicate the 75th and 25th percentile, and whiskers show the 95th and 5th percentile. Data is combined from 4 individual experiments with means weighted by the inverse of the variance of each replicate for (A1) **p-value <0.001, *’ p-value=0.054 with n *rg1-1* = 58, n *rg2-2* = 54 and n Col-0 = 65, for (A2) *p-value = 0.027 with n *rg1-1* = 58, n *rg2-2* = 54 and n Col-0 = 65. **B).** PVX virulence assay on TRV-VIGS *N. benthamiana* plants silenced for RanGAP2, RanGAP1 or combinations thereof in *N. benthamiana*. Silenced plants were infiltrated at 21 days post TRV-VIGS treatment with Agrobacteria for expression of the amplicon of either PVX105 or PVX106. Infiltrated leaf samples were harvested at 3 dpi for viral quantification by DAS-ELISA. Statistically significant difference was compared to TRV:GFP samples using the Wilcoxon-Signed Rank test (α = 0.05) with n = 24 for all samples represented. *p-value <0.05 and **p-value <0.001.

Similarly, we tested the role of RanGAP1/RanGAP2 in cyst nematode infection. As TRV-VIGS of RanGAP1/2 in tomato and potato roots appeared to be inefficient (**Supplemental Fig. S1)**, we took advantage of an alternative plant system to test the contribution of RanGAP2 and RanGAP1 to cyst nematode parasitism. For this, the *Arabidopsis thaliana* mutants *rangap1* (*rg1-1*) and *rangap2* (*rg2-2*) (23) were challenged with the beet cyst nematode *Heterodera schachtii*, which has a similar mode of parasitism as *G. pallida* on potato. Our data indicate that the total number of nematodes infecting the roots of *rg1-1* was significantly lower as compared to the wild-type control (Col-0), after 2 weeks of infection **(Fig. 2A1)**. Although this is less significant in *rg2-2*, a consistently lower trend was observed between different experimental repeats. In cyst nematodes, sex determination is dependent on environmental conditions. Auspicious conditions favour the development of female over male nematodes. Therefore, we also investigated the ratio between male and female nematodes at 2 weeks post-infection. Interestingly, both *rg1-1* and *rg2-2* plants harbour significantly fewer females than wild-type plants (**Fig. 2A2**). The reduction in the total number of nematodes and ratio of females infecting the roots of mutant plants collectively pinpoint that both RanGAP homologs contribute to the susceptibility of the roots of *A. thaliana* and thus, to cyst nematode virulence. Combined with the disease assay performed in *N. benthamiana*, we demonstrate that RanGAP1/2 contribute to the infection of roots and shoots by cyst nematodes and PVX, respectively.

### Gp-RBP-1 and PVX-CP associate with the RanGAP2-WPP domain *in planta*

To gain further insight into the effector targeting of RanGAP2, we next resolved the RanGAP domains involved in the association with Gp-RBP-1 and PVX-CP. Plant RanGAPs are characterized by a unique, N-terminal WPP domain (so-called for a conserved Tryp-Pro-Pro motif), which anchors the protein to the nuclear envelope (17). To test if the WPP domain is sufficient for the interaction with *G. pallida* and PVX effectors, we co-expressed GFP/CFP-tagged versions of the effectors with a RanGAP2-WPP variant tagged with the red fluorescent protein mCherry and a nuclear localization signal (NLS). The NLS-tagged RanGAP2-WPP was targeted to localize exclusively in the nucleus. On the other hand, both Gp-RBP-1 and PVX-CP have a more or less equal nucleocytoplasmic distribution (24, 25). It was anticipated that co-expressing WPP-NLS-mCh would shift the subcellular localization of these effectors towards the nucleus when these proteins exist in the same complex. This shift in nucleocytoplasmic distribution can be quantified by determining the fluorescence intensity ratio between the GFP-tagged protein in the nucleus and cytoplasm (I_N_/I_C_), as described previously (23). Confocal imaging was performed at 2 days post infiltration (2 dpi). Free CFP, which does not form a complex with the RanGAP2-WPP construct, was used as a negative control. Remarkably, our imaging data show that higher fluorescent intensities ratios for Rook4-GFP-4×HA, D383-1-GFP-4×HA, CFP-CP106, and CFP-CP105 occurred during co-expression with WPP-NLS-mCh in support of an interaction (**Fig. 3**). Apparently, the WPP domain of RanGAP2 is sufficient for complex formation with Gp-RBP-1 and PVX-CP. The nucleocytoplasmic distribution of the CFP negative control was not altered when co-expressed with the WPP-NLS-mCh construct. Overall, our findings demonstrate that the association of Gp-RBP-1 and PVX-CP locates to the WPP domain in RanGAP2.

**Fig. 3.**
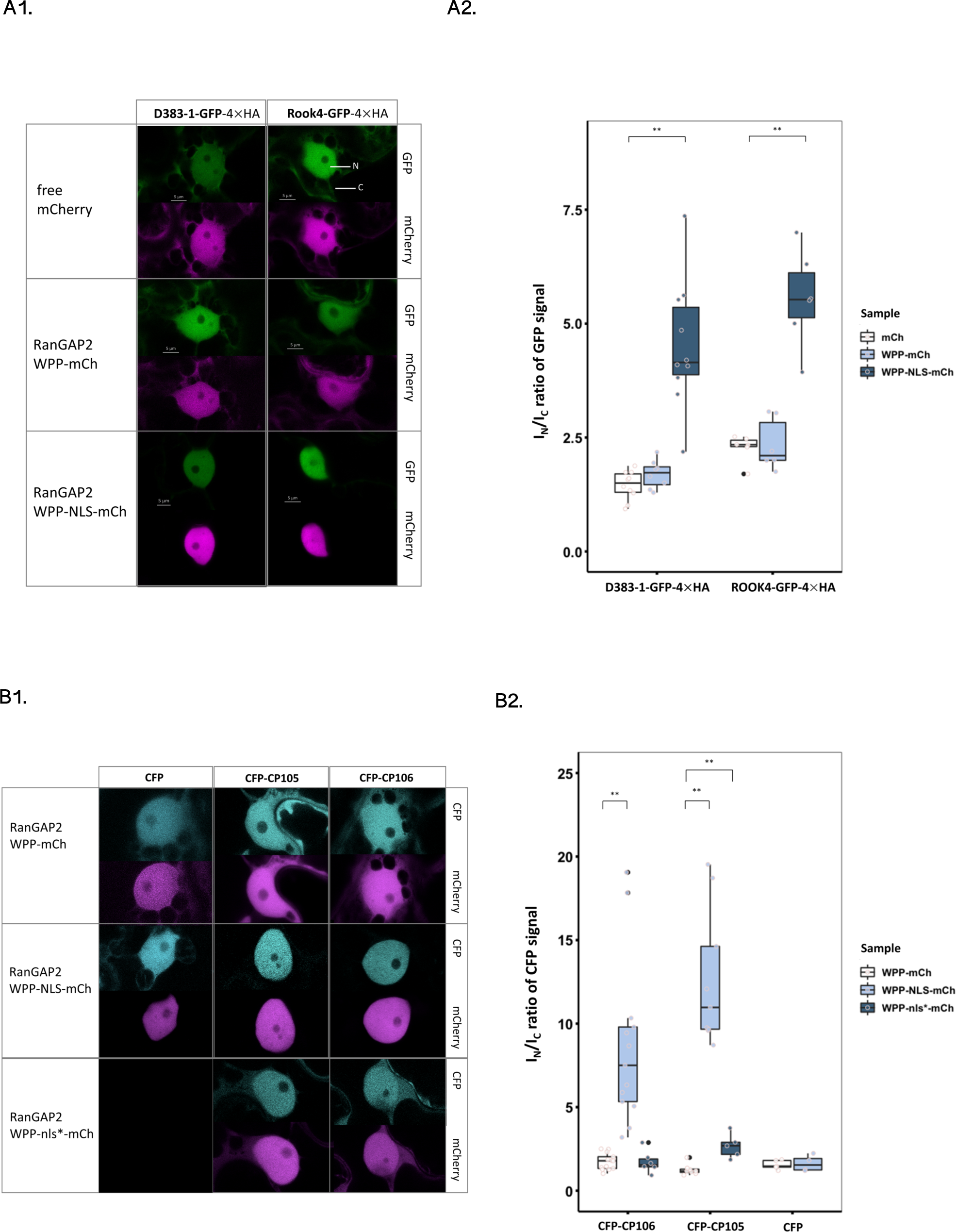
The WPP domain of RanGAP2 is sufficient for the interaction with Gp-RBP-1 (A) and PVX-CP (B). Representative confocal images of nuclei (N) and surrounding cytoplasm (C) of cells expressing mCherry-tagged RanGAP2-WPP and GFP-tagged Gp-RBP-1 (Rook4 or D383-1) (**A1**) or CFP-tagged PVX-CP constructs (PVX-CP 106 or 105) (**B1**). The CFP/GFP and mCherry channels are shown side by side for each combination. Quantification of the fluorescence intensity ratios (I_N_/I_C_ ratio) is represented in the accompanying boxplots (**A2** and **B2**). Boxes indicate the interquartile range with whiskers indicating the maximum and minimum values. Statistical significance difference was calculated using the Wilcoxon-Signed Ranked test with α = 0.05 with *p-value <0.05 and **p-value <0.001. For both PVX-CP and Gp-RBP-1, data shown is the combination of at least two independent repeats.

### Effector targeting of RanGAP2 does not affect the RanGAP2-receptor complex

RanGAP2 forms a heteromeric complex with Gpa2 and Rx1 *in planta*, which relies on an interaction between the receptor CC domain and the RanGAP2-WPP region (15, 16, 26). Our data indicate that Gp-RBP-1 and PVX-CP also interact at the WPP domain. It is, therefore, conceivable that effector targeting could affect the assembly of the RanGAP2-receptor complex. To explore this, full-length Gpa2 N-terminally tagged with 4×Myc was co-expressed with RanGAP2-GFP and either D383-1-8×HA or Rook4-8×HA. The coat proteins 4×HA-CP106 and 4×HA-CP105 were also included as controls as they do not activate Gpa2 but bind RanGAP2. Agroinfiltrated leaves were harvested at 2 dpi, before a visible cell death response occurred and enabled sufficient protein for detection by Western blot. Indeed, immunoblotting showed that co-expressing these effectors did not strongly affect the protein levels of 4×Myc-Gpa2 and RanGAP2-GFP at this time point (**Fig. 4A**). To study the interaction between Gpa2 and RanGAP2 under influence of the co-expressed effectors, we performed a Co-IP with 4×Myc-Gpa2 as bait and RanGAP2-GFP as prey. RanGAP2-GFP was pulled down with 4×Myc-Gpa2, but not in the absence of 4×Myc-Gpa2 as bait. Co-expressing with either Gp-RBP-1 or PVX-CPs did not alter the amount of RanGAP2-GFP pulled down along with 4×Myc-Gpa2 (**Fig. 4A**). Therefore, the interaction between Gpa2 and RanGAP2 is not affected by the cell death eliciting variant of Gp-RBP-1 at 2 dpi.

**Fig. 4.**
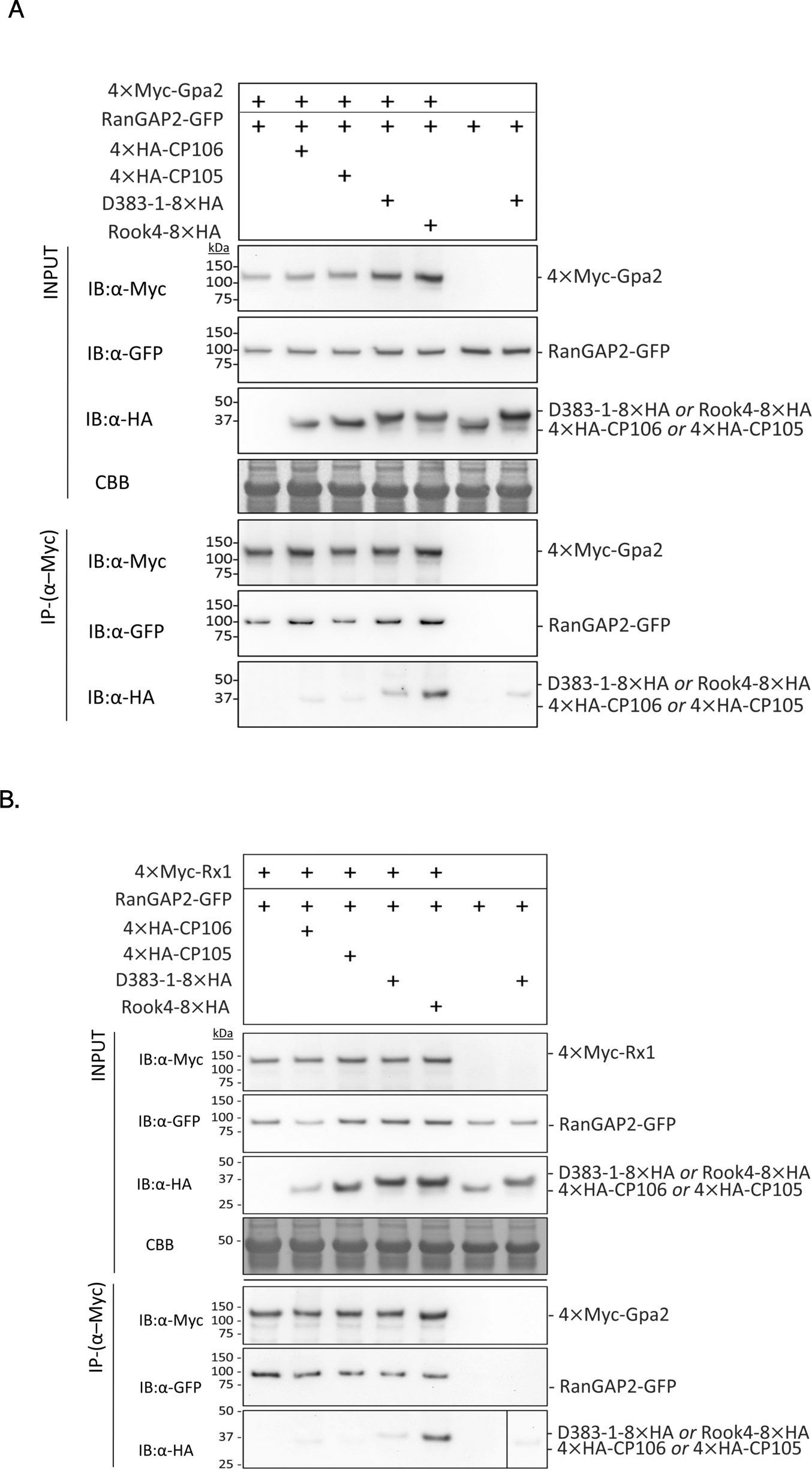
Effector targeting of RanGAP2 does not hamper its association with Gpa2 and Rx1. Shown in the figure are immunoblots from Co-IP experiments where 4×Myc-tagged Gpa2 or Rx1 constructs were used as bait and RanGAP2-GFP as prey. HA-tagged versions of CP106, CP105, or the Gp-RBP-1 D383-1 and Rook4 were additionally co-expressed. The top three immunoblot (IB) panels represent the input material. Coomassie brilliant blue (CBB) stained blots on which RuBisCO is visible are used as a control for equal loading for the input material. The lower three panels show the protein pulled down in the α-Myc immunoprecipitation. **A).** Co-immunoprecipitation to test if the interaction between full-length Gpa2 and RanGAP2 is affected by the eliciting and non-eliciting PVX-CPs or by the Gp-RBPs D383-1 or Rook4. The samples were harvested at 48 hours post agroinfiltration before cell death would occur in the combination of Gpa2 and D383-1. **B).** Co-immunoprecipitation to test if the interaction between full-length Rx1 and RanGAP2 is affected by the same sets of effectors as in (**A**). The proteins were co-expressed for 24 hours and harvested before cell death would occur in the combination of Rx1 and CP106. Data shown is representative of three independent repeats. “+” indicates the presence of a construct in the infiltration combination. Coomassie brilliant blue (CBB) stained blots serve as a loading control for the input material.

We likewise investigated whether the complex of Rx1/RanGAP2 would be affected by its interaction with PVX-CP. To test this, 4×Myc-Rx1 was co-expressed with RanGAP2 and the PVX-CPs for 24 hours before the leaves were harvested. At this time point, no cell death was visible, and the protein levels were sufficient for detection on Western blot. Immunoblotting shows that Rx1 and RanGAP2-GFP protein levels were not affected by the co-expressed effectors. Following IP using α-Myc beads, RanGAP2-GFP specifically co-immunoprecipitated with 4×Myc-Rx1, but not with α-Myc beads alone (**Fig. 4B**). The presence of the effectors does not interfere with the complex between Rx1 and RanGAP2. These findings are in agreement with that observed for Gpa2. Overall, we, therefore, conclude that the association between Gpa2 and Rx1 with RanGAP2 appears unchanged by the cell death eliciting effectors at this time-point. We corroborated these findings by examining the impact of effector targeting on the Rx1-CC and RanGAP2 interaction alone, which does not elicit cell death. Our immunoprecipitation data shows that PVX-CP and Gp-RBP-1 also do not affect complex formation between Rx1-CC with the RanGAP2 (**Supplementals Fig. S2**). Together, these data suggest that neither the eliciting nor non-eliciting effector variants induced the dissociation of the CC for complex formation with RanGAP2.

## DISCUSSION

The RanGAP2 protein has long been established as a co-factor of the closely-related intracellular NB-LRR immune receptors Gpa2 and Rx1 (15, 16). Nevertheless, how RanGAP2 functions in immunity provided by Gpa2 and Rx1 remains unclear. In this study, we expand further on this function by providing evidence for the physical association of RanGAP1 and RanGAP2 with the corresponding effectors of Gpa2 and Rx1, namely Gp-RBP-1 and PVX-CP. Our interaction data suggest that complex formation of effector variants of Gp-RBP-1 and PVX-CP with RanGAP1 and RanGAP2 is independent of the recognition specificity of Rx1 and Gpa2. Moreover, both RanGAP homologs contribute to infection of roots and shoots by these pathogens, a role not yet reported for these key host cell components. These data combined suggest that RanGAP2 and RanGAP1 are common host targets of evolutionary distinct effectors from two unrelated plant pathogens with entirely different life strategies.

Our finding that RanGAP2 and RanGAP1 are common host targets of unrelated pathogens is in line with an emerging picture that diverse pathogens utilize a small and overlapping set of host proteins to benefit their fitness (27). The PVX-CP and Gp-RBP-1 effectors bear no sequence or structural resemblance. Nevertheless, we show that both effectors can form a heteromeric complex with RanGAP2 *in planta* (**Fig. 1A** and **1B**). The convergence of multiple, unrelated effector molecules on a single host protein is proposed to facilitate pathogens to shift to new hosts and/or effectively suppress ‘defence hubs’ (22). Prominent examples being the molecular chaperone EDS1 and protease Rcr3 (28, 29). These common targets may fulfil a key function in a limited range of cellular processes that pathogens require for survival (27). This pinpoints that a fundamental cellular process regulated by RanGAP1 and RanGAP2 is targeted to facilitate disease progression. This, in turn, aligns with our findings demonstrating that the targeting of RanGAPs is observed for distinct pathogens in diverse plant backgrounds.

To our knowledge, this is the first report for the role of RanGAP1 and RanGAP2 in pathogenicity. Remarkably, a greater effect of the depletion of RanGAP1 was consistently found during infection by cyst nematodes in *A. thaliana* as compared to RanGAP2. This suggests that the RanGAP2 and RanGAP1 homologs may have a yet undefined, differing role in processes that cyst nematodes exploit for their fitness. Notably, Gp-RBP-1 was found to pull-down less efficiently with RanGAP1 (**Fig. 1A**). Immunoblotting assays indicate that both RanGAP homologs are expressed at comparable levels, minimizing the likelihood that the observed quantitative differences in interaction are due to RanGAP1 being present in lower abundance. Alternatively, the difference in interaction between RanGAP1 and RanGAP2 is likely caused by intrinsic differences between the RanGAP2 and RanGAP1 proteins, with the *N. benthamiana* homologs sharing an overall sequence identity of 66.29% (65.26% at the WPP domain and 67.61% at the LRR domain). However, it remains to be determined whether the differences observed in disease assays are linked to this sequence variation or binding affinities of RanGAP1 and RanGAP2 with Gp-RPB-1.

During interphase, RanGAP is involved in the maintenance of a RanGTP/RanGDP gradient required for macromolecule transport between the plant nucleus and the cytoplasm (17). Interestingly, both nucleocytoplasmic trafficking, as well as mitotic activity, are crucial host cell processes involved in nematode and viral pathogenicity and cyst nematodes (30, 31). Cyst nematodes establish a feeding site inside the host roots, which acts as a nutrient sink to support the growth and reproduction of the nematode (32). The formation of such a feeding structure also called syncytium involves drastic molecular and metabolic changes of the root cell, including the reactivation of the cell cycle and the incorporation of neighbouring cells via progressive cell wall dissolution (33). Hence, nematodes may recruit RanGAP1 and RanGAP2 to modulate cellular processes involved in syncytium formation. Interestingly, previous sequence analysis reveals Gp-RBP-1 to harbour high homology to Ran-binding protein, RanBPM (9), further supporting the hypothesis that cyst nematodes may target RanGAP1 and RanGAP2 to modulate the Ran cycle for their own benefit.

Like cyst nematodes plant viruses fully depend on their host cells for replication and spreading disease, which could explain why they share a common host target despite a different mode of action. PVX is a positive-stranded RNA virus whereby the cytoplasm is their primary site of replication (34). Nonetheless, there is accumulating evidence for the interplay between plant RNA viruses and the plant nucleus. In line with this, several viral proteins having been described to translocate to the nucleus for functions such as suppressing RNAi and/or recruit for splicing factors necessary to modulate viral mRNA (35, 36). Interestingly, at least one virus is known to disrupt this gradient by targeting Ran to interfere with nuclear efflux of antiviral factors (37). Thus, nucleocytoplasmic trafficking may constitute an important aspect of plant-RNA virus infection. However, whether targeting by PVX-CP directly impacts these RanGAP2-related functions and the precise implications thereafter require more concrete molecular and biochemical studies.

We further demonstrate that the interaction of both Gp-RBP-1 and PVX-CP locate to the WPP domain of RanGAP1/2 (**Fig. 3**). The WPP domain is characteristic of a small family of proteins associated to the nuclear envelope and possibly exclusive to plants (reviewed in (38)). This domain mediates, together with WPP-interacting proteins (WIPs) and WPP-interacting tail-anchored proteins, the localisation of the RanGAPs to the outer surface of the nuclear envelope (NE) during interphase (18, 39, 40). Targeting of the WPP may thus collectively disturb the cellular distribution and GAP activity of RanGAP2, affecting the overall biological functions of the protein (e.g., in nuclear trafficking). It would, therefore, be interesting to see how PVX and cyst nematodes benefit from interacting with RanGAPs as a common virulence target during host infection.

In addition, RanGAP2 is also a co-factor of the potato immune receptor Gpa2 and the observed direct interaction between Gp-RBP-1 and RanGAP2 *in planta* is in accordance with the effects of artificial tethering of Gp-RBP-1 and RanGap2 described by (Sacco *el al.*, 2009). The direct interaction of RanGAP2 with Gp-RBP-1 supports the idea that plant RanGAP2 may play a role in mediating the indirect recognition of the effector by Gpa2 (15, 16). Moreover, the role of RanGAP2 as a cytoplasmic retention factor also coincides with previous findings that Rx1 needs to be localized in the cytoplasm for recognition (24). The physical association of RanGAP2 with the corresponding effectors of its immune regulators reported here further reinforces this model. It is worth noting that previous studies could not establish the complex formation of RanGAP2 with PVX-CP by Co-IP (15). We attribute these differences to variation in platforms and setups used. Most notably, earlier approaches made use of C-terminally tagged PVX-CP constructs (19). Here, PVX-CP tagged at the N-terminus was employed instead as the C-terminal variant has been proven to compromise viral infection (25). Whether the loss of CP function is directly linked to impaired RanGAP2 binding also warrants further investigation.

Based on our data, we propose RanGAP2 could serve as a bait that facilitates direct effector recognition (41). We, therefore, propose that recognition of PVX-CP/Gp-RBP-1 is a two-step event according to the bait-and-switch model (41). This model involves the initial ‘docking’ of the effectors to the bait, in this case, RanGAP2 via the WPP domain. However, the landing of effectors to RanGAP2 is insufficient for recognition and subsequent receptor activation, given that the non-eliciting effectors also bind. Instead, docking to RanGAP2 brings the effector in closer proximity to the LRR, which is then able to directly sense structural determinants on an accessible/exposed side of the effector. Such a model is in accordance with the divergence of the Rx1 and Gpa2 LRRs to sense structurally unrelated effectors. It is also in line with our Co-IP experiments showing that RanGAP2, Rx1/Gpa2, and their matching effectors may possibly exist as a concurrent formation of a tripartite complex (**Fig. 5**). The function we ascribe for RanGAP2 is reminiscent of that described for the extra Solanaceous Domain (SD) in the Sw-5b receptor protein, which is likewise postulated to enhance effector detection by the LRR when there is low amounts of the effector present (42). The attributed role of RanGAP2 could be further linked to the finding that the N-terminus of Gp-RBP-1 mediates the binding to RanGAP2 with variation in this region having been described to contribute to the strength in inducing Gpa2-mediated HR (9) Specifically, tagging Gp-RBP-1 at the N-terminus with a fluorophore prevents energy transfer in FRET-FLIM assay and co-localization with a WPP-NLS construct (**Supplementals Fig. S4**). This suggests that the N-terminus may be involved in RanGAP2 binding. On the other hand, a proline to serine substitution at position 187 in the C-terminus determines recognition specificity but is not required for the binding of RanGAP (9). This polarization in regions of Gp-RBP-1 required for RanGAP2 binding and recognition further reinforces the function of RanGAP2 as a molecular bait in the NLR switch model as proposed (ref).

**Fig. 5.**
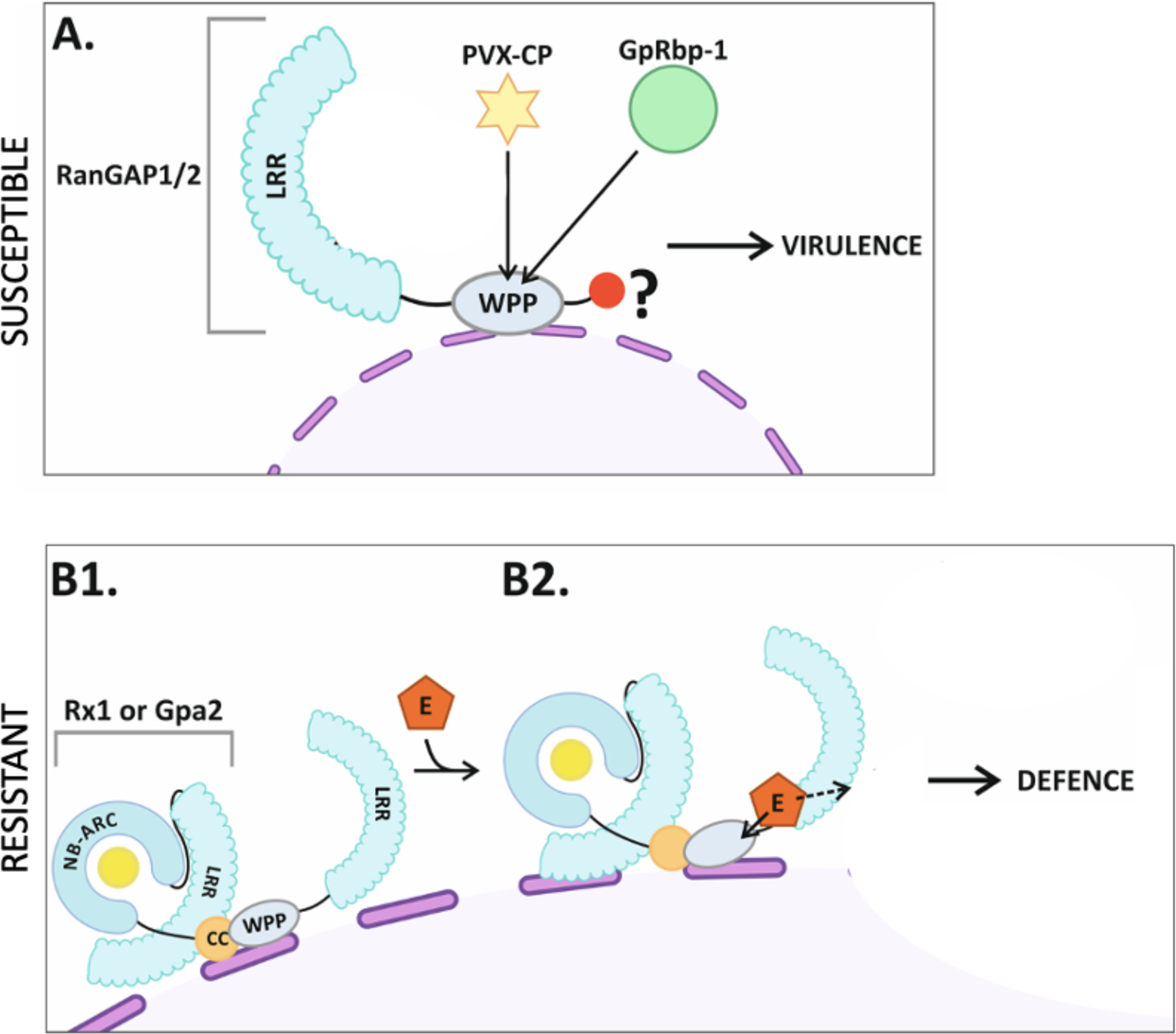
Working model for the dual role of RanGAP1/2 in pathogen virulence (A) and recognition events by Gpa2/Rx1 (B). In the latter case, a model for only RanGAP2 is shown, for which a clearer role in resistance of Gpa2 and Rx1 has been established. **A).** In the absence of a matching NB-LRR receptor, Gp-RBP-1 and PVX-CP effectors target both RanGAP1 and RanGAP2 via the WPP to promote viral disease and nematode feeding cell formation in a yet undisclosed manner (**red circle**). **B**). When Rx1/Gpa2 is present, the receptors tether to RanGAP1 and RanGAP2 at the nuclear envelope via its WPP domain (**B1**). During invasion, both eliciting and non-eliciting forms of Gp-RBP-1 and PVX-CP (orange pentagon labelled ‘E’) initially dock to RanGAP2, bringing the effector in close proximity to the immune receptor (**B2**) to facilitate recognition. Successful recognition leads to downstream events towards defence.

Alternatively, RanGAP2 could act as a effector target guarded by two NB-LRRs with distinct recognition specificities. In this model, we anticipate that effector targeting of RanGAP2 would perturb RanGAP1/2 and indirectly trigger recognition. However, we were unable to detect any apparent changes in the stability, size, or banding pattern of RanGAP1, RanGAP2 or its bound receptors in our assays. We do not rule out the possibility that the effectors may impose other or more subtle modifications leading to the perturbations of RanGAP1/2, which does not involve the dissociation of the heteromeric complex. The role of RanGAP2 as a guardee, however, contradicts earlier works detailing the lack of positively-selected residues on the RanGAP2 surface (43). This is expected from a guarded host protein as it would need to co-evolve with the pathogen. The future challenge therefore lies in uncovering the exact molecular basis of RanGAP2-mediated activation of Rx1/Gpa2-like immune receptors by specific effector variants. Our finding that Gp-RBP-1 and PVX-CP interact with RanGAP homologs provides an important stepping-stone towards this goal.

## MATERIALS AND METHODS

### Plasmid constructs

To obtain Gp-RBP-1 variants D383-1 and Rook4 with N or C-terminally tagged GFP or HA, the target genes were initially subcloned into the pRAP vector by NcoI/KpnI digestion (44). A similar strategy was followed for cloning of CFP-tagged PVX-CPs. For transient expression experiments in *N. benthamiana*, the tagged-effector constructs were finally subcloned into the pBIN+ binary vector and transformed into *Agrobacterium tumefaciens* strain MOG10 (6).

Heterologous expression by *A. tumefaciens* transient assay in *N. benthamiana*

Heterologous protein expression was carried out by *A. tumefaciens* transient assay (ATTA) in plants, as described previously (24). Briefly, Agrobacteria strains carrying the expression vectors were grown in Yeast Extract Broth (YEB) medium (5 g/L peptone, 1 g/L yeast extract, 5 g/L beef extract, 5 g/L sucrose and 2.5 g/L NaCl and 2 ml 1M MgSO_4_) overnight. Grown bacterial cells were spun down and re-suspended in infiltration medium and optical densities at wavelength 600 nm (OD_600_) were adjusted to final OD_600_ values of 0.2-0.4 for all constructs in co-immunoprecipitation and imaging assays unless otherwise stated. *A. tumefaciens* suspensions were then infiltrated on the abaxial surface of the leaves of *N. benthamiana* plants using needleless syringes. Infiltrated spots were harvested for protein extraction or examined by microscopy at 2 days post infiltration (dpi).

### Transient silencing by TRV-VIGS and PVX resistance assay in *N. bethamiana*

Constructs used for RanGAP1, RanGAP2 and RanGAP1 + 2 TRV-VIGS silencing in *N. bethamiana* are as described previously (15, 19). Agroinfiltration was performed in a similar way as for the *N. benthamiana* agroinfiltrations (see above). Briefly, bacteria are grown overnight in YEB medium and re-suspended in MMAi containing 200 μM acetosyringone. Final OD_600_ of TRV1 and TRV:Rg1, TRV:RG1+2 or TRV:RG2 mix were adjusted to 0.5 for infiltration. Infiltrated plants were grown for 21 days to allow for systematic silencing before use in viral infection assays as described previously ((45)). Briefly, Agrobacteria carrying amplicons for PVX105 or PVX106 were infiltrated on TRV-silenced plants at OD_600_ values of 0.002. Between 1-5 dpi, 13 mm leaf discs were harvested from infiltrated spots, extracted in phosphate buffer (pH = 7) and finely ground using Tissuelyzer II (Qiagen) with settings of 30 seconds at 30 Hz. Ground materials were incubated in a 96-well plate coated with polyclonal antibody targeted against the PVX-CP (Prime Diagnostics) at 37°C for 2 hours, before a second round of incubation with a conjugate antibody carrying alkaline phosphatase. Viral levels were quantified by absorbance measurements at 405 nm with the BioRad microplate reader (model 680) following a reaction with the substrate *p*-Nitrophenol. Statistical analyses were performed in R studio Version 1.1.456. Data from assays performed in this study were checked for normality using Shapiro-Wilk Test. Depending upon the outcome of the normality test, statistical level was determined either by T-test or Wilcoxon-Signed Rank Test with α = 0.05 as specified in the text.

### Virus induced gene silencing in potato or tomato

Constructs used for RanGAP1 and RanGAP2 VIGS silencing in potato and tomato are described previously (15, 19). Agroinfiltration was performed in a similar way as for the *N. benthamiana* agroinfiltrations (see above). Briefly, bacteria are grown overnight in YEB medium and re-suspended in MMA containing 200µM acetosyringone. Final ODs of a TRV1 and TRV:Rg1.1, TRV:Rg1.2, TRV:RG1+2 or TRV:RG2 mix are adjusted to 0.3 for infiltration in potato and to 0.4 for infiltration in tomato. Potato and tomato plants are grown and maintained in silver sand under standard greenhouse conditions. For nematode infection approximately 1000 eggs or 12.000 eggs of *G. pallida* (Rookmaker) were added to the potato or tomato plants, respectively. Relative gene expression was calculated with the ΔΔCt method (46)with RPN7 (47). For tomato, normalization was done using the geometric mean of reference genes tubulin (48) and MST1.

### *In planta* co-immunoprecipitation and detection of recombinant proteins

Total protein extracts were prepared by grinding leaf material in protein extraction buffer (20% (v/v) glycerol, 50 mM Tris-HCl pH 7.5, 2 mM EDTA, 300 mM NaCl, 0.6 mg/ml Pefabloc SC plus (Roche, Basel, Switzerland), 2% (w/v) polyclar-AT polyvinylpolypyrrolidone (Serva, Heidelberg, Germany), 10 mM dithiothreitol and 0.1% (v/v) Tween20) on ice. For co-immunoprecipitation, protein extracts were passed through a Sephadex G-25 column (GE Healthcare, Chicago, Illinois) and pre-cleared by treatment with rabbit-IgG agarose (Sigma, 50 μL slurry per 60 μL protein extract). The cleared protein extract was incubated with μMACS α-GFP paramagnetic (Miltenyi, Bergisch Gladbach, Germany) for 1h at 4°C. Columns were washed with washing buffer (20% (v/v) glycerol, 50 mM Tris-HCl pH 7.5, 2 mM EDTA, 300 mM NaCl, 0.10% (v/v) Nonidet 40 and 5mM dithiothreitol) five times and eluted by removing the column from the uMACS collector and adding 45uL of the washing with the washing solution. The input samples were mixed with 1X NuPage LDS sample buffer with 0.25 M dithiothreitol and incubated at 95°C for 5 minutes.

For Western blotting, proteins were separated by SDS-PAGE on NuPage 12% Bis-Tris gels (Invitrogen) and blotted to 0.45 μm polyvinylidene difluoride membrane (Thermo Scientific). Before immunodetection we blocked the membranes for 1h at room temperature in 5% (w/v) powder milk in PBS with 0.1% Tween20. For immunodetection rabbit α-GFP (Abcam, Cambridge, United Kingdom) with horseradish peroxidase-conjugated donkey α-rabbit (Jackson ImmunoResearch, Ely, United Kingdom) or horseradish peroxidase-conjugated rat α- HA (Roche) were used. Peroxidase activity was visualized using SuperSignal West Femto or Dura substrate (Thermo Scientific) and imaging of the luminescence with G:BOX gel documentation system (Syngene, United Kingdom).

### Confocal laser scanning and FRET-FLIM microscopy

Confocal microscopy was performed on *N. benthamiana* epidermal cells using a Zeiss LSM 510 confocal microscope (Carl-Zeiss) with a 40X 1.2 numerical aperture water-corrected objective. For co-localization studies the argon laser was used to excite at 488 nm for GFP and chlorophyll, and the HeNe laser at 543nm to excite mCherry. GFP and chlorophyll emission were detected through a band-pass filter of 505 to 530nm and through a 650 nm long-pass filter, respectively. mCherry emission was detected through a band-pass filter of 600 to 650nm. Nuclear and cytoplasmic fluorescence intensities were quantified using ImageJ (49). For FRET-FLIM analysis, the FRET between GFP and mCherry was detected via Fluorescent Lifetime Imaging Microscopy. The HYD SMD detector of a Leica SP5 CLSM (Leica, Wetzlar, Germany) was used to measure the emission and fluorescent lifetime of GFP (495-545 nm) and the red fluorescent mCh emission (570-625 nm). The excitation of the GFP chromophore was measured using a white light laser (488 nm). The Time-correlated single-photon counting (TCSPC) was performed using a Becker & Hickl FLIM system FLIM analysis of TCSPC was performed with the B&H SPCImage software (Becker & Hickl GmbH, Berlin, Germany).

### Nematode infection assays in *A. thaliana*

*rangap1-1* (SALK_058630) and *rangap2-2* (SALK_006398) seeds were obtained from the Nottingham Arabidopsis Stock Centre (23). All *A. thaliana* genotypes used in the experiments are in the Columbia 0 (Col-0) genetic background. The presence of T-DNA inserts in the lines was confirmed by PCR using specific primers designed with the iSect Primers tool of the SIGNAL SALK database (**Supplementals Table S1**), in combination with the universal LB primer (50). For nematode assays, seeds were vapour sterilized and vernalized at 4°C in the dark for 4 days to break seed dormancy. After vernalisation the seeds were plated in pairs in 9cm petri dishes containing modified KNOP medium. Plants were grown at 25°C under a 16h/8h light-dark cycle. 10 day-old seedlings were inoculated with 60-70 surface-sterilized *H. schachtii* infective juveniles. After 2 weeks of infection, the number of males and females present in the roots of Arabidopsis plants were counted visually and the size of females and syncytia were calculated with Leica M165C Binocular (Leica Microsystems, Wetzlar, Germany) and the Leica Application Suite software (Leica Microsystems). To combine results from 4 biological replicates, we weighted the measures of association from each replicate by the inverses of their variances. The variance of such weighted average is simply the inverse of the sum of the inverses of the variances which allow standard methods to be used to test for the overall significance at the 5% level of the genotype and the number of nematodes per plant. Such approach corresponds to methods to combine studies under a fixed effects model.

## ACKNOWLEDGEMENTS

The works described in this study benefits from the financial support of the COST Action SUSTAIN FA1208 as well as Dutch Top Technology Institute Green Genetics (5CFD051RP), Dutch Technology Hotel grant, the Dutch Technology Foundation STW and Earth and Life Sciences ALW (STW-GG 14529), NWO project 828.11.002, and TTI Green Genetics grant 4CC058RP which are part of the Netherlands Organization for Scientific Research (NWO).We are thankful to Dr. Jan Willem Borst (Laboratory of Biochemistry, Wageningen University & Research) and the microspectroscopy research facility (Wageningen University & Research) for access to the imaging equipments and their technical expertise. We also thank Dr. Matthieu Joosten (Laboratory of Phytopathology, Wageningen University & Research) for providing WPP-NLS, WPP-nls*, RanGAP1-GFP, RanGAP2-GFP and RanGAP2-mCherry constructs.

## AUTHOR CONTRIBUTIONS

Conceptualization A.G.; Methodology, O.C.A.S., and A.D.G.M; Investigation, O.C.A.S., A.D.G.M, E.J.S., H.O., C.S., S.P., J.R., R.P., and A.E.; Writing – Original Draft, O.C.A.S. and A.D.G.M; Writing – Review & Editing, O.C.A.S., A.D.G.M., E.J.S., A.G., and G.S; Funding Acquisition, A.G., and G.S.

## SUPPORTING INFORMATION CAPTIONS

**Supplemental Table S1.** Identity percentage of RanGAP2 and RanGAP1 sequences from potato, tomato and *N. benthamiana*

**Supplemental Fig. S1 Inefficient silencing of RanGAP1/2 in roots of tomato** (***Solanum lycopersicum***) **and potato** (***Solanum tuberosum***). Tobacco rattle virus (TRV) carrying guide DNA fragments targeting RanGAP1 (RG1_a and RG1_b), RanGAP2(RG2), both homologues (RG1+2) or green fluorescent protein as negative control was inoculated to the leaves of 10-day old tomato seedlings to induce transient virus-induced silencing of RanGAP1/2. **A)** RanGAP1 or **B)** RanGAP2 expression was measured by quantitative RT-PCR in TRV-infected plants and compared to the expression on TRV-GFP-infected plants. Values are normalized to the geometric mean of reference genes tubulin (Aimé et al., 2013) and MST1. Individual samples are composed of ∼10 plants/construct and RT-PCR measurements are performed in triplicate. TRV-mediated silencing was quantified 3 weeks after inoculation in the leaves of inoculated plants. Silencing was efficient for RanGAP1 using construct RG1_a **C)** 3 weeks after TRV infection plants were inoculated with ∼12000 eggs of *G. pallida* (Rookmaker) and were grown for 2 months to allow completion of the nematode life cycle. After 2 months of nematode inoculation, cysts were extracted and counted from the complete root systems of plants with efficient RanGAP1 silencing. **D)** a similar set-up was used for TRV-mediated transient silencing in potato, with inoculum being ∼1000 eggs. Cysts present in the roots of VIGS-potato were extracted and quantified and no difference was found between mean amount of cysts present in potatoes inoculated with RG1, RG2, RG1/2 and GFP-silencing TRV.

**Supplemental Fig. S2 Supporting Figure 3.** Expression of **A)** *RanGAP2* and **B)** *RanGAP2* in the roots of *H. schachtii*-inoculated *Arabidopsis*, after 2, 7, 10 and 14 days of inoculation. Expression compared to mock-inoculated plants was determined by quantitative RT-PCR. The relative expression of RanGAP1 and RanGAP2 was normalised to the geometric mean of reference genes Ubiquitin 5 (Anwer et al., 2018) and ubiquitin carboxyl-terminal hydrolase 22 (Hofmann & Grundler, 2007). **C)** Size of female nematodes and syncytia established in the roots of *rg1-1* and *rg2-2*, with Col-0 as wild-type control. Sizes are shown in mm^2^. Data from 4 biological repeats is combined, with means weighted by the inverse of the variance of each biological repeat. Stars indicate a significant difference as established by a linear fit, * p-value= 0.015 with n*_rg1-1_* = 109, n*_rg2-2_* = 80 and n_Col-0_= 129

**Supplemental Fig. S3 The binding of RanGAP2 to the CC domain of Rx1 is not disrupted in the presence of the effectors studied.** Co-immunoprecipitation investigating whether the interaction between the CC domain of Rx1 and RanGAP2 is affected by the coat proteins of non-eliciting and eliciting PVX-CP strains or by the Gp-RBPs D383-1 or Rook4. The samples were harvested at 48 hours post agroinfiltration. As a control for aspecific binding, 4×Myc-GFP was used as bait. “+” indicates the presence of a construct in the co-expressed combination.

**Supplemental Fig. S4. Tagging Gp-RBP-1 with a fluorescence protein at the N-terminus impairs its interaction with RanGAP2 (WPP domain). A).** Confocal imaging of RanGAP2-WPP-NLS constructs co-expressed with Rook4 or D383-1 tagged with GFP at the N or C termini. Representative images of nuclei in infiltrated *N. benthamiana* epidermal cells for combinations involving N-terminally tagged Gp-RBP-1s are given in **A1**. Key: N = nucleus; C = cytoplasm. Quantification of cellular distribution by I_N_/I_C_ measurements is summarized in boxplot of **A2** with boxes representing the interquartile range. Data shown is the combination of two experimental repeats. **B).** Boxplot indicating lifetime (picoseconds) from a FRET-FLIM experiment whereby full length RanGAP2-WPP-mCh is co-expressed with the same set of effectors as described in **A2**. Data shown is pooled from three experimental repeats.

**Supplemental Fig. S5.** PVX virulence assay on TRV-VIGS *N. benthamiana* plants silenced for RanGAP2 in *N. benthamiana*. Silenced plants were infiltrated at 21 days post TRV-VIGS treatment with Agrobacteria for expression of the amplicon of either PVX105 or PVX106. Infiltrated leaf samples were harvested at 5 dpi for viral quantification by DAS-ELISA. Statistically significant difference was compared to TRV:GFP samples using the Wilcoxon-Signed Rank test (α = 0.05) with n = 8 for all samples represented.

**Supplemental Table S2.** List of primers used in the study for the genotyping of *A. thaliana* RanGAP mutants.

