## Supporting Information for "Two evolutionary distinct effectors from a nematode and virus target RanGAP1 and 2 via the WPP domain to promote disease"

The following Supporting Information is available for this article:

**Supplemental Table S1.** Identity percentage of RanGAP2 and RanGAP1 sequences from potato, tomato and *N. benthamiana*

| ***RanGAP1*** | | | | |
|---|---|---|---|---|
| % | ***S. tuberosum*** | ***S. lycopersicum*** | ***N. benthamiana*** | ***A. thaliana*** |
| ***S. tuberosum*** | | 97.944 | 92.150 | 69.951 |
| ***S. lycopersicum*** | 97.944 | | 84.857 | 70.195 |
| ***N. benthamiana*** | 92.150 | 84.857 | | 69.343 |
| ***A. thaliana*** | 69.951 | 70.195 | 69.343 | |
| ***RanGAP2*** | | | | |
| % | ***S. tuberosum*** | ***S. lycopersicum*** | ***N. benthamiana*** | ***A. thaliana*** |
| ***S. tuberosum*** | | 95.683 | 90.499 | 67.764 |
| ***S. lycopersicum*** | 95.683 | | 86.121 | 67.827 |
| ***N. benthamiana*** | 90.499 | 86.121 | | 67.643 |
| ***A. thaliana*** | 67.764 | 67.827 | 67.643 | |

**Supplemental Fig. S1 Poor silencing of RanGAP1/2 in tomato (*Solanum lycopersicum*) and potato (*Solanum tuberosum*)**. Tobacco rattle virus (TRV) carrying guide DNA fragments targeting RanGAP1 (RG1_a and RG1_b), RanGAP2(RG2), both homologues (RG1+2) or green fluorescent protein as negative control was inoculated to the leaves of 10-day old tomato seedlings to induce transient virus-induced silencing of RanGAP1/2. **A)** RanGAP1 or **B)**RanGAP2 expression was measured by quantitative RT-PCR in TRV-infected plants and compared to the expression on TRV-GFP-infected plants. Values are normalized to the geometric mean of reference genes tubulin (Aimé et al., 2013) and MST1. Individual samples are composed of ~10 plants/construct and RT-PCR measurements are performed in triplicate. TRV-mediated silencing was quantified 3 weeks after inoculation in the leaves of inoculated plants. Silencing was efficient for RanGAP1 using construct RG1_a **C)** 3 weeks after TRV infection plants were inoculated with ~12000 eggs of *G. pallida* (Rookmaker) and were grown for 2 months to allow completion of the nematode life cycle. After 2 months of nematode inoculation, cysts were extracted and counted from the complete root systems of plants with efficient RanGAP1 silencing. **D1** and **D2)** a similar set-up was used for TRV-mediated transient silencing in potato, with inoculum being ~1000 eggs. Cysts present in the roots of VIGS-potato were extracted and quantified and no difference was found between mean amount of cysts present in potatoes inoculated with one of two constructs VIGS constructs targeting RanGAP1 (RG1_a or RG1_b), with both RanGAP1-targeting constructs (RG1_a + RG1_b), and with a construct targeting GFP. Similarly, no difference was found in the amount of cysts present in potatoes inoculated with constructs targeting RanGAP2 or GFP.

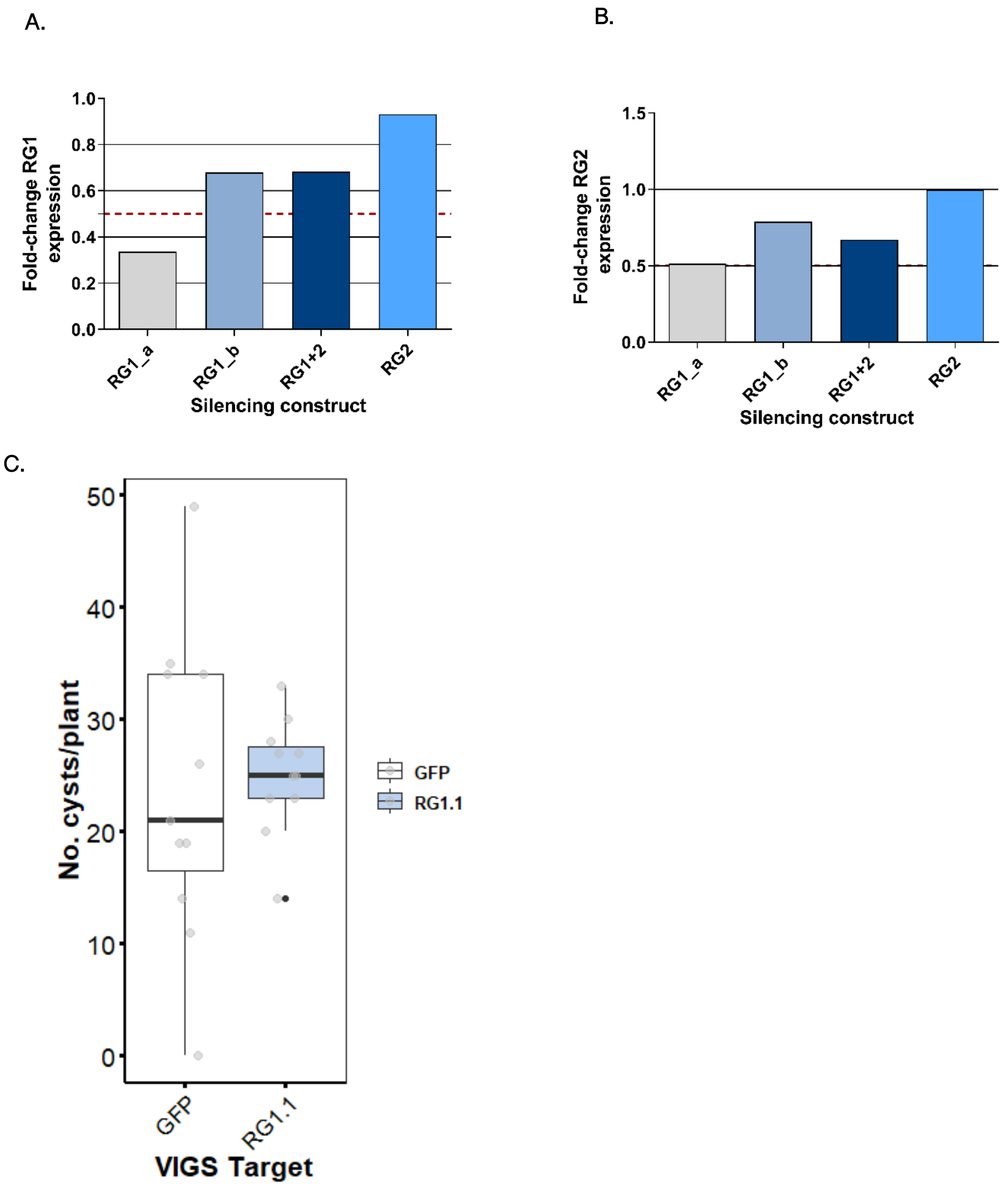
A.
Fold-change RG1 expression
1.0
0.8
0.6
0.4
0.2
0.0
RG1_a
RG1_b
RG1+2
RG2
Silencing construct
B.
Fold-change RG2 expression
1.5
1.0
0.5
0.0
RG1_a
RG1_b
RG1+2
RG2
Silencing construct
C.
No. cysts/plant
50
40
30
20
10
0
GFP
RG1.1
VIGS Target
GFP
RG1.1

D1.

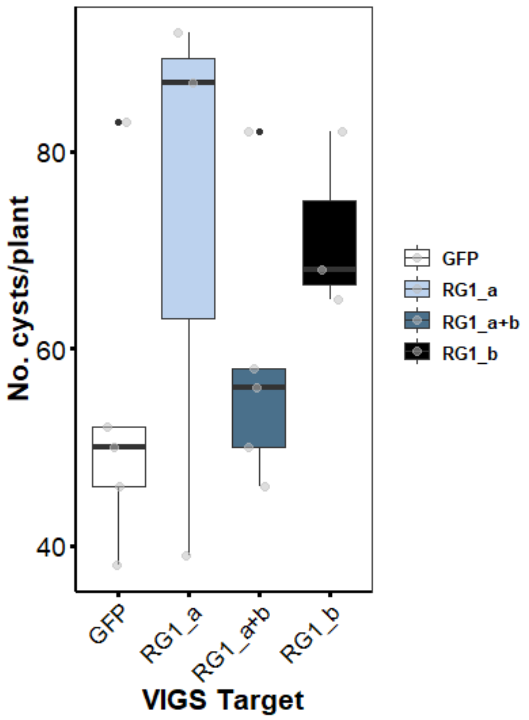

No. cysts/plant
80
60
40
GFP
RG1_a
RG1_a+b
RG1_b
VIGS Target
GFP
RG1_a
RG1_a+b
RG1_b


D2.

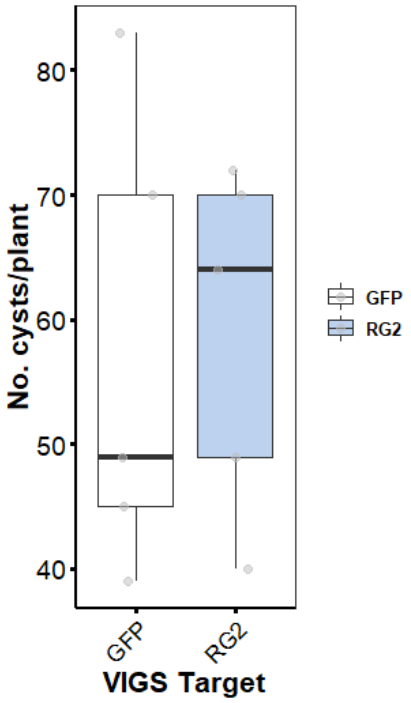

No. cysts/plant
80
70
60
50
40
GFP
RG2
VIGS Target
GFP
RG2

**Supplemental Fig. S2.** Expression of **A)** *RanGAP2* and **B)** *RanGAP2* in the roots of *H.schachtii*-inoculated *Arabidopsis*, after 2, 7, 10 and 14 days of inoculation. Expression compared to mock-inoculated plants was determined by quantitative RT-PCR. The relative expression of RanGAP1 and RanGAP2 was normalised to the geometric mean of reference genes Ubiquitin 5 (Anwer et al., 2018) and ubiquitin carboxyl-terminal hydrolase 22 (Hofmann & Grundler, 2007). **C)** Size of female nematodes and syncytia established in the roots of *rg1-1* and *rg2-2*, with Col-0 as wild-type control. Sizes are shown in $mm^2$. Data from 4 biological repeats is combined, with means weighted by the inverse of the variance of each biological repeat. Stars indicate a significant difference as established by a linear fit, * p-value= 0.015 with $n_{rg1\text{-}1}$ = 109, $n_{rg2\text{-}2}$ = 80 and $n_{Col\text{-}0}$= 129

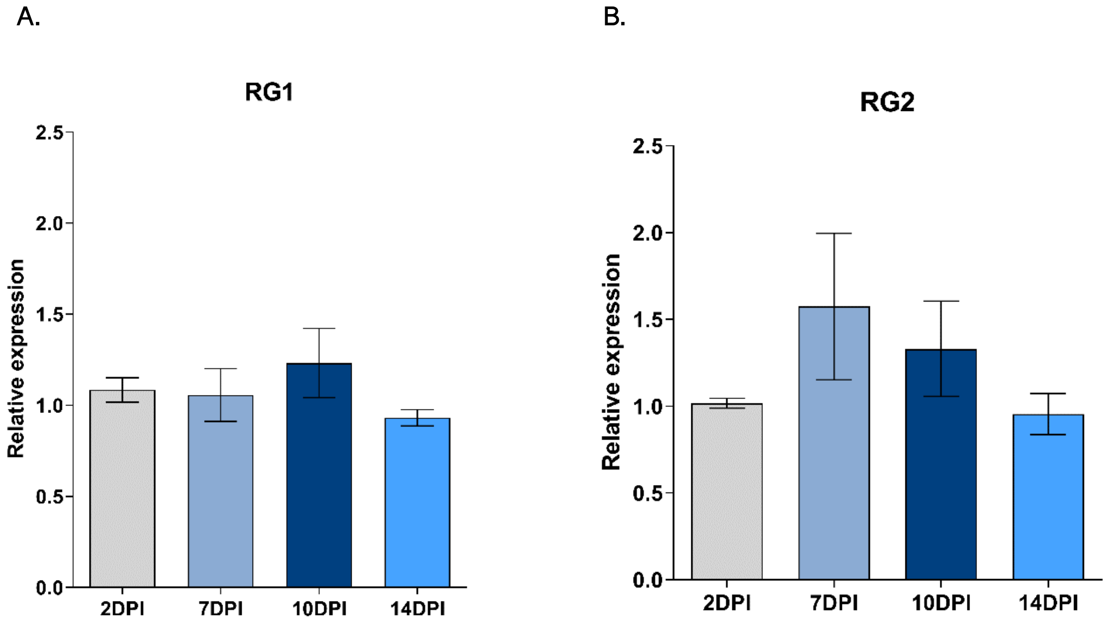

C.

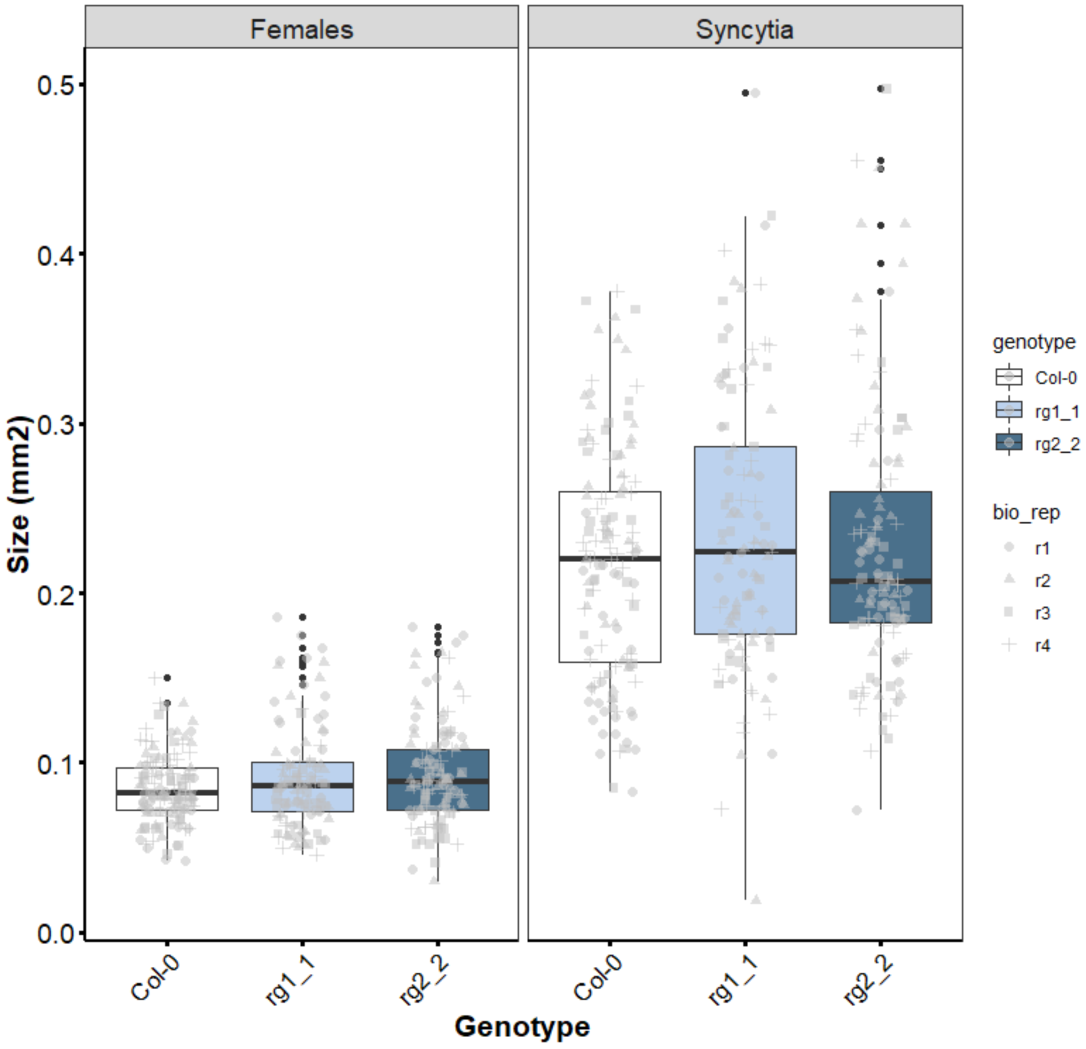

Females
Syncytia
0.5
0.4
0.3
0.2
0.1
0.0
Size (mm2)
Col-0
rg1_1
rg2_2
Col-0
rg1_1
rg2_2
Genotype
genotype
Col-0
rg1_1
rg2_2
bio_rep
r1
r2
r3
r4

**Supplemental Fig. S3 The binding of RanGAP2 to the CC domain of Rx1 is not disrupted in the presence of the effectors studied.** Co-immunoprecipitation investigating whether the interaction between the CC domain of Rx1 and RanGAP2 is affected by the coat proteins of non-eliciting and eliciting PVX-CP strains or by the Gp-RBPs D383-1 or Rook4. The samples were harvested at 48 hours post agroinfiltration. As a control for aspecific binding, 4×Myc-GFP was used as bait. "+" indicates the presence of a construct in the co-expressed combination.

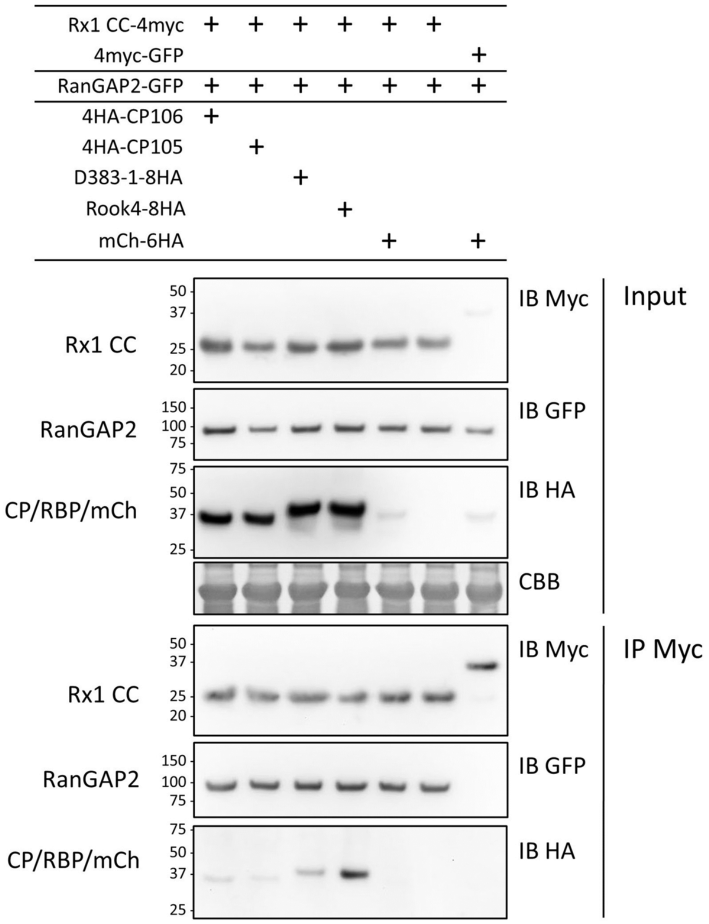

**Supplemental Fig. S4. Tagging Gp-RBP-1 with a fluorescence protein at the N-terminus impairs its interaction with RanGAP2 (WPP domain). A).** Confocal imaging of RanGAP2-WPP-NLS constructs co-expressed with Rook4 or D383-1 tagged with GFP at the N or C termini. Representative images of nuclei in infiltrated *N. benthamiana* epidermal cells for combinations involving N-terminally tagged Gp-RBP-1s are given in **A1**. Key: N = nucleus; C = cytoplasm. Quantification of cellular distribution by $I_N/I_C$ measurements is summarized in boxplot of **A2** with boxes representing the interquartile range. Data shown is the combination of two experimental repeats. **B).** Boxplot indicating lifetime (picoseconds) from a FRET-FLIM experiment whereby full length RanGAP2-WPP-mCh is co-expressed with the same set of effectors as described in **A2**. Data shown is pooled from three experimental repeats.

A1.

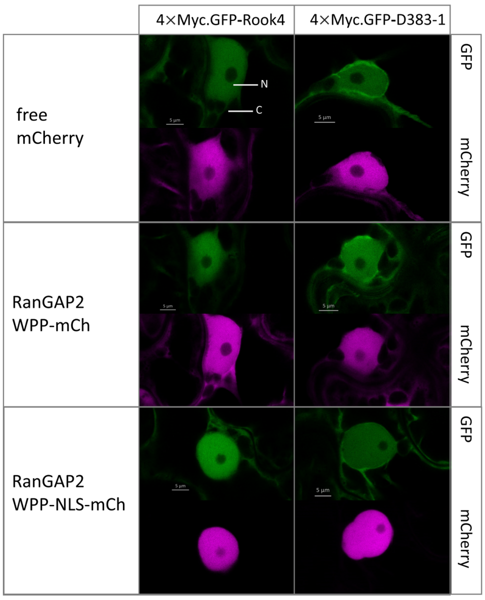


A2.

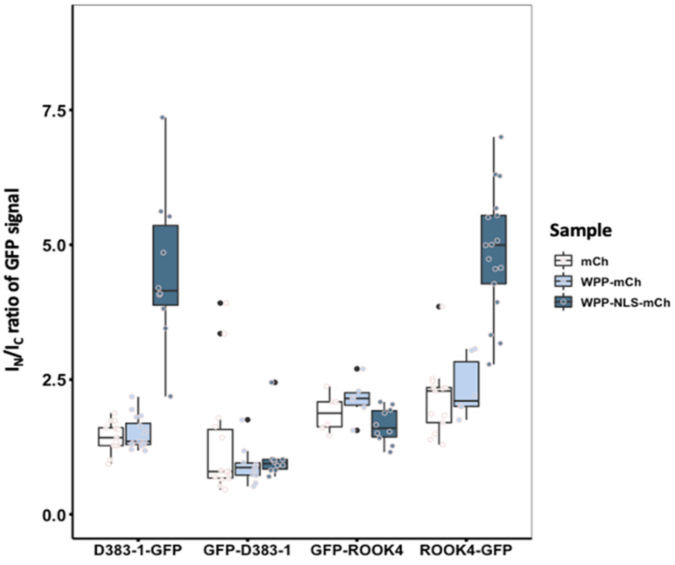

B.

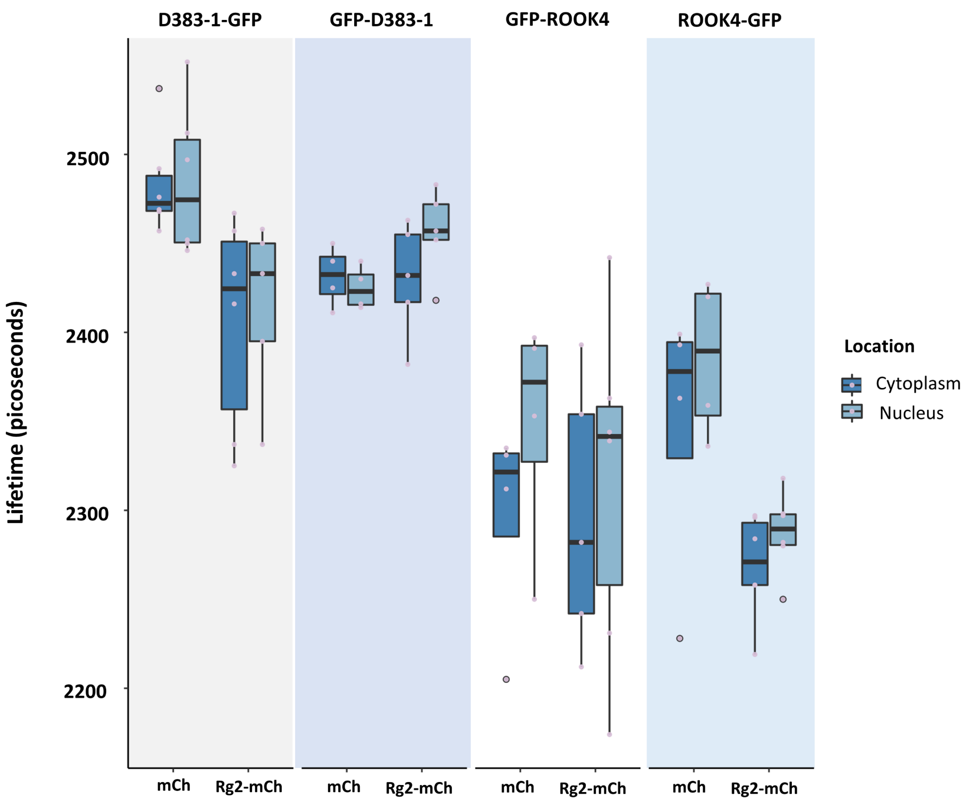
D383-1-GFP
GFP-D383-1
GFP-ROOK4
ROOK4-GFP
Lifetime (picoseconds)
2500
2400
2300
2200
mCh
Rg2-mCh
mCh
Rg2-mCh
mCh
Rg2-mCh
mCh
Rg2-mCh
Location
Cytoplasm
Nucleus

**Supplemental Fig. S5.** PVX virulence assay on TRV-VIGS *N. benthamiana* plants silenced for RanGAP2 in *N. benthamiana*. Silenced plants were infiltrated at 21 days post TRV-VIGS treatment with Agrobacteria for expression of the amplicon of either PVX105 or PVX106. Infiltrated leaf samples were harvested at 5 dpi for viral quantification by DAS-ELISA. Statistically significant difference was compared to TRV:GFP samples using the Wilcoxon-Signed Rank test (α = 0.05) with n = 8 for all samples represented.

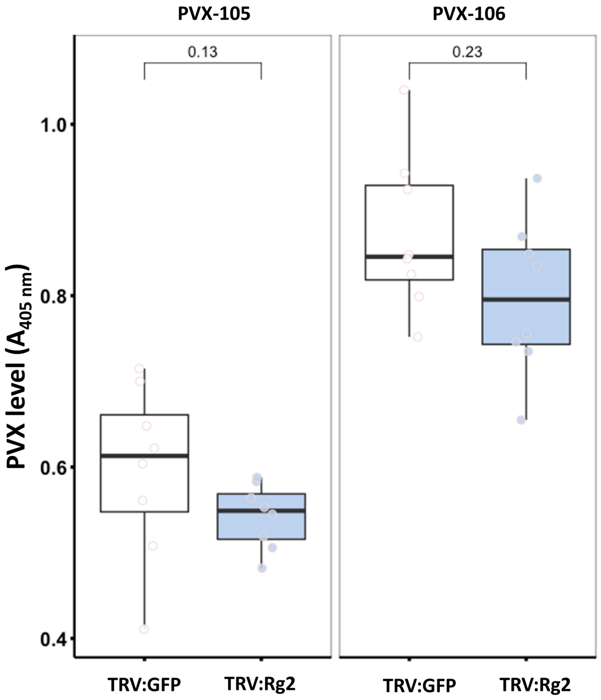

**Supplemental Table S2.** List of primers used in the study for the genotyping of *A. thaliana* RanGAP mutants.

| *Primer name* | *Sequence* |
|---|---|
| SALK_058630_LP (*rg1-1*) | 5'- CAAAGGAACAAGCTTGTCCAG-'3 |
| SALK_058630_RP (*rg1-1*) | 5'- TCCATTTCTCCAACGAATCTG- '3 |
| SALK_006398_LP (*rg2-2*) | 5'- CTCCTCTGATCACAATCGGTC- '3 |
| SALK_006398_RP (*rg2-2*) | 5'- TGGTCTTTGGTTAAGCTACCG –'3 |
